# BitBIRCH Clustering Refinement Strategies

**DOI:** 10.1101/2025.03.20.644337

**Authors:** Kenneth López Pérez, Kate Huddleston, Vicky Jung, Ramón Alain Miranda-Quintana

## Abstract

Chemical libraries are becoming not only increasingly bigger, but they are doing so at an accelerated pace. Keeping up with this explosion in chemical data demands more than just hardware upgrades, we need dramatically more efficient algorithms as well. We have been working in this direction, with the introduction of the iSIM framework, which uses *n*-ary similarity to speed up the processing of very large sets. Recently, we showed how to use this technique to cluster billions of molecules with unprecedented efficiency through the BitBIRCH algorithm. In this Application Note we present a package fully-dedicated to expanding on the BitBIRCH method, including multiple options that give the user appreciable control over the tree structure, while dramatically improving the quality of the final partitions. Remarkably, this is achieved without compromising the efficiency of the original method. We also present new post-processing tools that help dissect the clustering information, as well as ample examples showcasing the new functionalities. BitBIRCH is publicly available at: https://github.com/mqcomplab/bitbirch.

## 1. INTRODUCTION

The similarity principle, “*similar molecules have similar properties*”^1–3^, is at the core of the medicinal chemistry and cheminformatics fields. An obvious key part of this old dictum is the molecular similarity, which is usually defined over pairs of molecules, and commonly calculated with the Tanimoto^4,5^ similarity index using binary fingerprint representations^6,7^. Getting to know which molecules are similar to potentially active is key in the drug discovery and development process.^8–13^ Clustering (a classic unsupervised learning technique) can be critical in organizing data/molecules in groups that are similar or have related properties.^14,15^ Having well-resolved groups of compounds is key to good property prediction^16,17^, speeding virtual screening^18^, and identifying novel regions of chemical space^19,20^, to name a few instances.

The Taylor-Butina^21,22^ algorithm remains arguably the most popular clustering algorithm in the drug design community. Its core idea is quite simple: at any given iteration, pick the molecule with most neighbors up to a pre-specified similarity threshold, and then that “seed” compound and the corresponding neighborhood will form a cluster. This simplicity, however, comes at a steep computational price, since a pairwise matrix of comparisons (usually, using the Tanimoto similarity) is required, which demands O(*N*^2^) time and memory resources. We recently proposed BitBIRCH^23^ as an alternative to this problem that, without sacrificing the final clusters’ quality, has a much more attractive O(*N*) scaling. Our method builds on the *n*-ary similarity formalism and uses a tree-inspired data type to process all the molecules. However, at its core, BitBIRCH also works by finding candidate centroids for highly-populated regions of chemical space, and assigning molecules to them based on their similarity to the available centers. Methods with this “radial” characteristic (see also RTC^24,25^ and eQual^26^ in the molecular dynamics community) tend to form an overly large first cluster (Figs. 1A and 1B) that can overlap with distant regions in chemical space (Fig. 1C).

**Figure 1:**
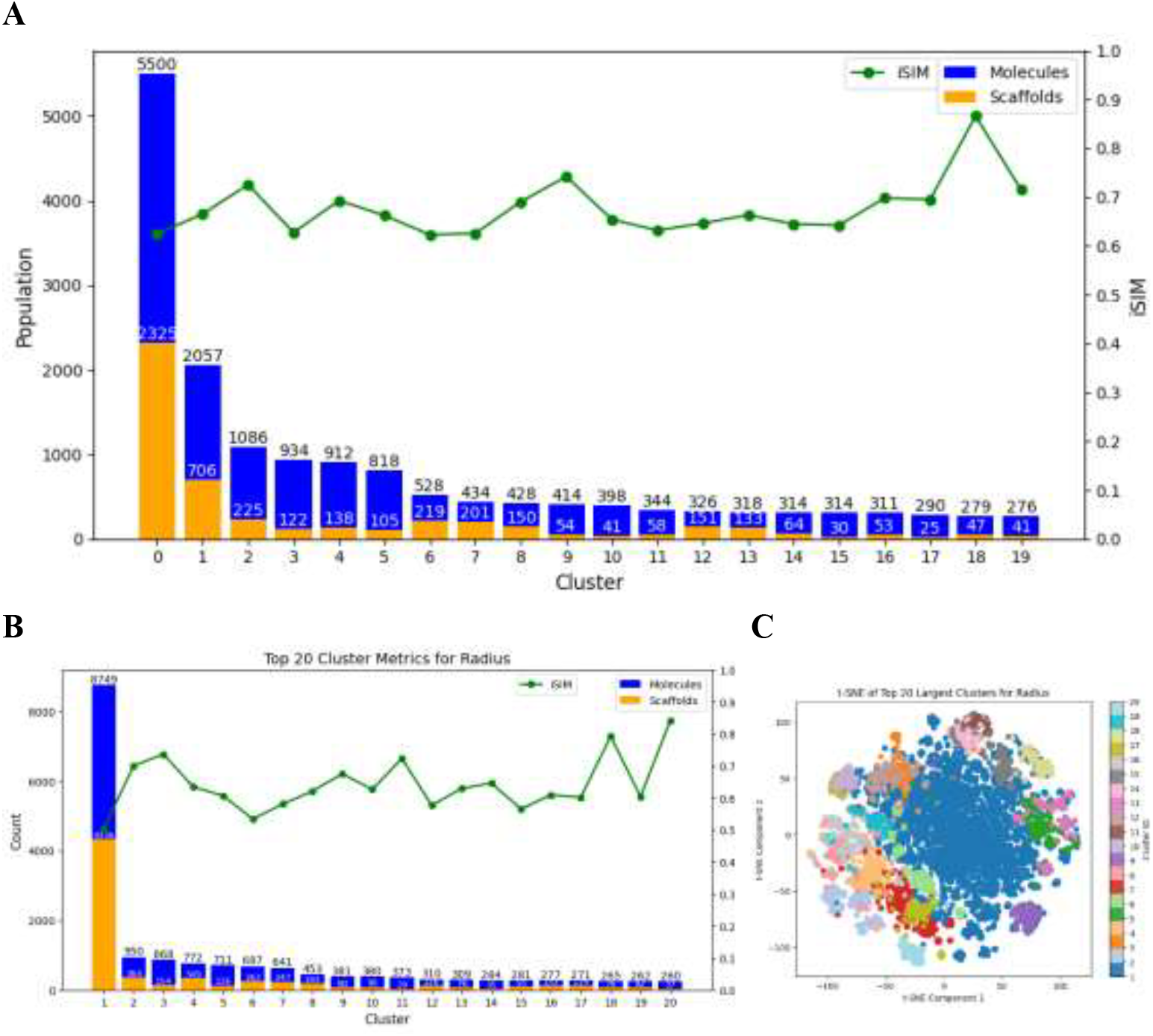
Analysis of the top 20 most populated clusters from the ChEMBL natural products set using the Taylor-Butina (A) and BitBIRCH with radius merge (B). Total number of molecules (blue, and left vertical axis), number of unique scaffolds (green), and iSIM values (red, and right vertical axis) for each cluster. (C) t-SNE projection of the top 20 most populated clusters from BitBIRCH with radius merge.

In this work, we not only introduce a fully-dedicated BitBIRCH package, but we also discuss several alternatives on how to improve the final clustering. After briefly introducing the basics of the iSIM^27^ and traditional BitBIRCH formalisms in the next section, we discuss how the resulting BitBIRCH tree can be manipulated to avoid the concentration of compounds in one large cluster. This greatly increases the flexibility of our package, and gives the user the tools to mold the final clustering to their particular needs. We present examples of the usage and results of these new techniques, stressing the fact that they do not introduce any appreciable overhead relative to the standard BitBIRCH implementation. This package is freely available at https://github.com/mqcomplab/bitbirch.

## 2. THEORY: iSIM and BitBIRCH

BitBIRCH’s efficiency is a result of two key ingredients: a tree structure that reduces the number of comparisons required to cluster the molecules (Fig. 2A), and a bit feature (BF) encoding of the cluster information. The latter is just a compact representation containing, for the *j*^th^ cluster **(X**^(*j*)^), its number of molecules, *N* _*j*_, the indices of the molecules, mols _*j*_, the cluster centroid, **c** _*j*_, and a vector containing the (linear) sum of the columns of the fingerprints belonging to the cluster, **ls** _*j*_. It is not necessary to store each individual fingerprint to be able to determine key properties like the cluster radius and diameter, since iSIM allows to calculate these using only *N* _*j*_ and **ls** _*j*_. More precisely, the iSIM Tanimoto of the *j*^th^ cluster, *i*T (**X**^(*j*)^), represents the average of the 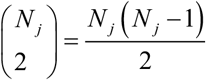 pairwise Tanimoto similarities between the molecules in the cluster:

**Figure 2:**
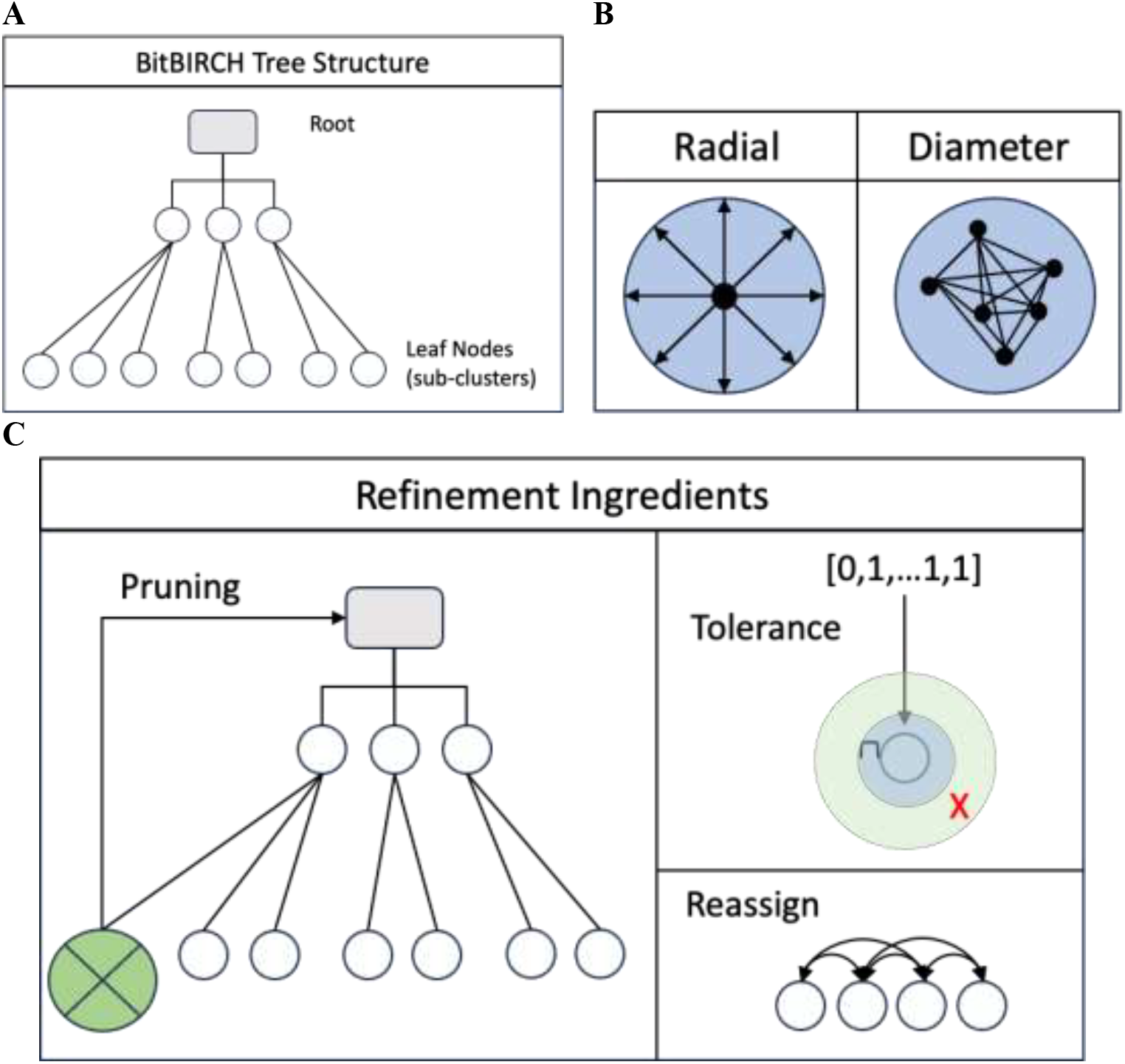
A: Schematic representation of a BitBIRCH tree, with root, non-leaf nodes, and leaf sub-clusters, B: Cartoon representation of the radial and diameter merge criteria, C: New functionality included in the BitBIRCH package: tree pruning, tolerance merge, and cluster reassignment.

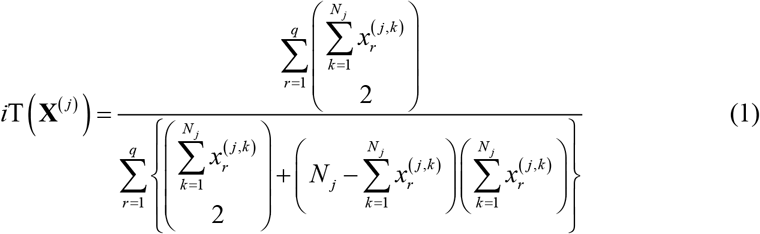

where 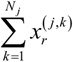 is the *r*^th^ component of **ls** _*j*_.

The cluster membership criterion in the original BitBIRCH implementation was based on the cluster radius, which can be calculated as:

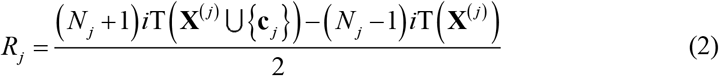

Notice that now we are explicitly considering a “similarity radius”, that is, the average Tanimoto similarity between all the molecules in the cluster and the cluster centroid, **c** _*j*_, which can be calculated as:

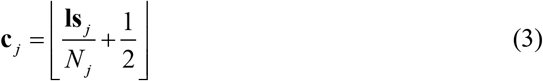

As noted in the Introduction, the radial criterion can result in an overly-populated top cluster. As expressed in Eq. (2), a new molecule will be added to a pre-existing cluster if it is similar enough to the centroid, but no direct connection to the other cluster members is enforced. Now, we propose a tighter merge criterion that explicitly monitors the relation between all the molecules in a cluster. For this, we use the traditional definition of the diameter of a cluster, *D*_*j*_, as the average of all the pairwise similarities of its elements.

However, this is just the iSIM of the cluster, so we simply need to calculate:

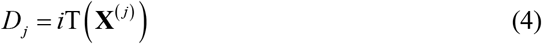

See Fig. 2B for a schematic representation of the difference between the radial and diameter cluster membership conditions.

As we will discuss in the forthcoming sections, the diameter condition results in tighter clusters, but there are still some cases in which the most populated cluster has a disproportionate amount of molecules and unique scaffolds. For this reason, our new BitBIRCH package contains three new options that greatly improve the final clustering results, without compromising the time and memory efficiency of this algorithm:

### Prune

Observations over multiple datasets have shown that the BitBIRCH clusters are extremely robust, with the possible exception of the most populated one. This is why we now introduce the pruning functionality which (as represented in Fig. 2C), removes a leaf cluster from the tree, updates the upper levels of the tree accordingly (adjusting the nodes’ centroids until the root level), and then reinserts those molecules into the tree again.

### Tolerance

When the new molecules are being reinserted into the tree, we need to guarantee that the quality of the remaining clusters is not affected. For this reason, we implemented a new cluster merge option that ensures that the following condition is satisfied when a new molecule, new_mol, is looking to be merged into the *j*^th^ cluster:

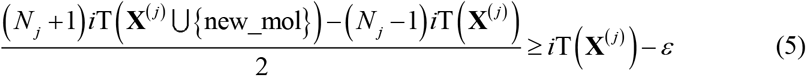

So, adding new_mol cannot decrease the diameter of the cluster in more than a pre-specified quantity, *ε*. For example, *ε* = 0 indicates that the new molecules are not allowed to decrease the average similarity between the elements of the set. It is up to the user to pick this parameter.

### Reassign

In the original BIRCH paper, the authors proposed a refinement protocol that effectively uses the tree only to identify potential centroids of high-density regions and then assigns the molecules to each of them. We include this functionality in this new version of BitBIRCH, where the user can pick any number of the most populated clusters (the default is 20), extract their centroids, and then screen the molecules from those clusters against the centroids and assign a compound to its closer centroid, as calculated using the Tanimoto similarity.

## 3. METHODOLOGY

### Data

To benchmark the new BitBIRCH functionalities, we used the ChEMBL33^28^ natural products database (n = 64,086). Molecules were represented with binary 2048-bit RDKit^29^ fingerprints.

### Refinement methods

Several functionalities were added to the BitBirch repository, including new cluster merging functions. A scheme with possible alternatives for merging functions and refinement method is shown in Figure 3.

**Figure 3:**
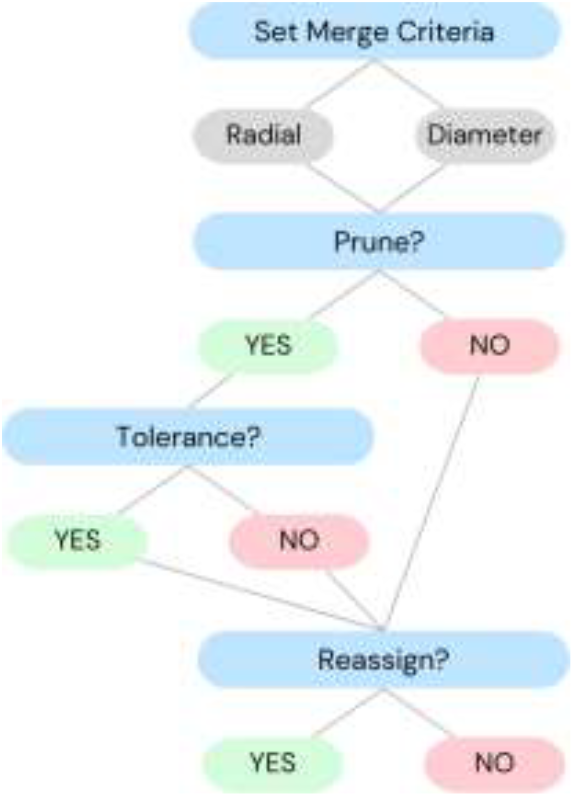
Flowchart indicating the different options in the new BitBIRCH package.

With the `set_merge(merging_criterion)` function, it is possible to set `‘radius’` or `‘diameter’` as the merging criterion to compare to the threshold. Another option for merging criterion is `‘tolerance’`, which uses a default of ε = 0.05 in Equation 5 to decide if subclusters/molecules should merge. Given the chance of having a big cluster, the `prune(fingerprints)` method prunes the most populated cluster from the tree and re-fits the corresponding molecules in the remaining structure of the three. For cases where similar molecules did not encounter each other because of the tree structure, we introduced the `reassign(`default: `top = 20)` method, which will compare the centroids of the most populated clusters and their molecules. Finally, we introduce `fit_BFs(BFs_list)`; this method will fit a new three starting from a list of BitFeatures Subclusters (`_BFSubcluster`), and not necessarily just individual molecules.

Implementations and examples of the presented methods are included in our GitHub repository: https://github.com/mqcomplab/bitbirch.

### Analysis

To characterize the clusters, we employed: the number of molecules, the number of unique Murcko^30^ scaffolds and iSIM (average similarity of the molecules in the set). The scaffolds were calculated using rdkit’s `Chem.Scaffolds.MurckoScaffold` module. t-SNE was used as a visualization for the clustering of the different method combinations. The timing analysis was done on the University of Florida’s HiPerGator supercomputer using one 10 GB node.

## 4. RESULTS

In Fig. 4 we show a detailed analysis of the different combinations of options in the BitBIRCH package. Note in Fig. 4A the immediate impact of changing the merging criterion from radius (with > 8,000 molecules and > 4,000 unique scaffolds in the most populated cluster) to diameter (with ∼3,000 molecules and ∼1,600 unique scaffolds, which are less than in the Taylor-Butina case, Fig. 1A). It is important to remark that all the diameter approaches directly control the diversity of the clusters, given that they are based on the iSIM values of the set. For example, using the radial criterion, the top cluster (Fig. 1B) has an average Tanimoto between pairs of ∼ 0.5, while the diameter merging guarantees that even the top cluster has an average Tanimoto above 0.65.

**Figure 4:**
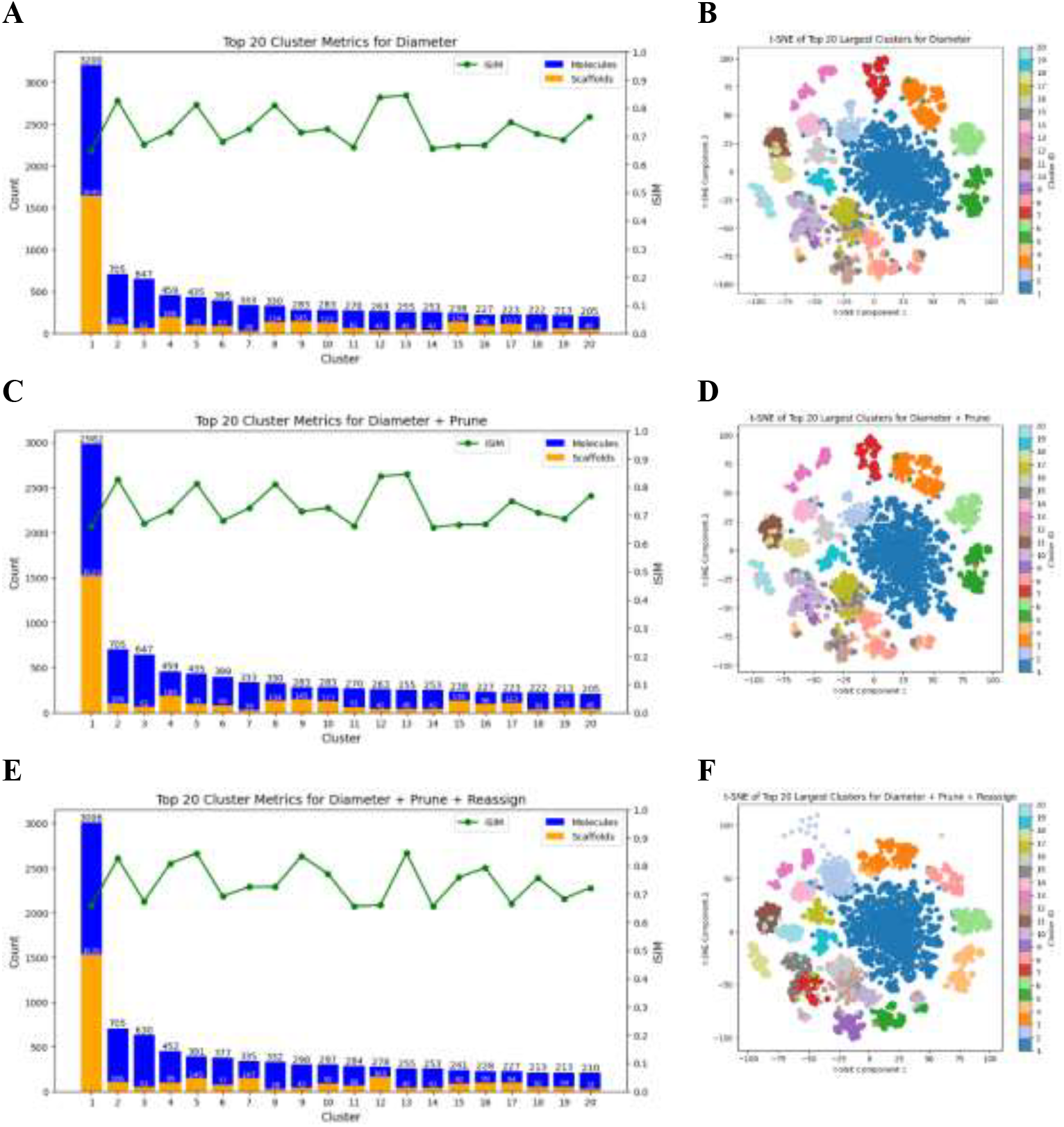
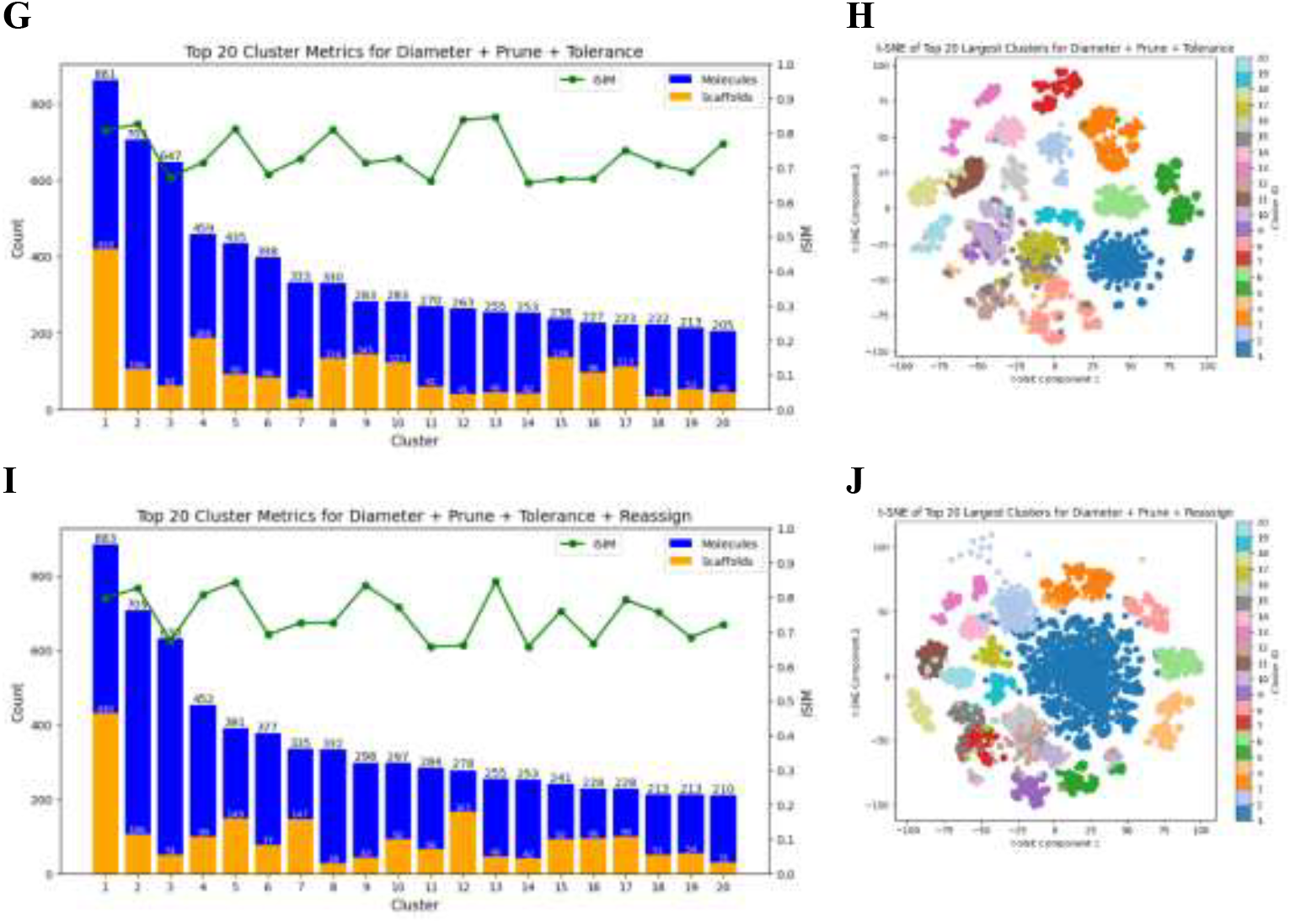
Analysis of the top 20 most populated clusters from the ChEMBL natural products set using the Diameter (A), Diameter + Prune (C), Diameter + Prune + Reassign (E), Diameter + Prune + Tolerance (F), and Diameter + Prune + Tolerance + Reassign (I) BitBIRCH variants. Total number of molecules (blue, left vertical axis), number of unique scaffolds (orange, left vertical axis), and iSIM values (green, and right vertical axis) for each cluster. t-SNE projection of the top 20 most populated clusters from (B), Diameter + Prune (D), Diameter + Prune + Reassign (F), Diameter + Prune + Tolerance (H), and Diameter + Prune + Tolerance + Reassign (J) BitBIRCH variants.

Diameter merge, while drastically improving over the formerly proposed radius merge, might still produce clusters that are too large (as seen in Fig. 4B) or that overlap different regions of chemical space. The prune option discussed above tries to correct this by passing the molecules of the top cluster through an updated tree, but as shown in Figs. 4C and 4D, simply pruning the tree does not drastically affect the distribution of the molecules. In the end, we still get ∼3,000 molecules in the top cluster, but now with a better distribution of scaffolds, at around 1,500. Although we pruned the most populated cluster and updated the remaining structure of the tree, the tree branching was initially influenced by the removed leaf. Reassigning the molecules of the top 20 clusters (Figs. 4E and 4F) keeps the pruned distribution almost identical, with only minor changes in the number of molecules and scaffolds and the overall distribution of the clusters in the 2D t-SNE representation.

The biggest improvement is seen with the tolerance options, which is no surprise since this guarantees that while the pruned molecules are added to the tree the remaining clusters’ quality is not negatively affected. Figs. 4G and 4I show that, with and without doing a final reassignment, the number of molecules in the top cluster drastically goes down to ∼900 molecules, which results in a much more uniform profile of populations per cluster. This is also reflected in a much-decreased count of unique scaffolds in the top clusters, which is another indicator of a much tighter partition. This is reflected in Figs. 4H and 4K have better separation in the projected chemical space.

Importantly (Table 1), the new BitBIRCH alternatives do not introduce a significant overhead on the previous implementation.

**Table 1:** Computing time comparison of the different BitBIRCH options over the ChEMBL33 natural products set.

| Method | Time (s) | $\Delta$ Time (s) |
| --- | --- | --- |
| Radial | 13.25 | 0.00 |
| Diameter | 13.31 | 0.06 |
| Diameter + Prune | 14.22 | 0.97 |
| Diameter + Prune + Tolerance | 14.28 | 1.03 |
| Diameter + Prune + Reassign | 14.30 | 1.04 |
| Diameter + Prune + Tolerance + Reassign | 14.27 | 1.02 |

Before concluding, we want to highlight that the tools described in sections 2 and 3 can be combined in several ways, which allows the BitBIRCH package to partition the data with great flexibility. For example, it is easy to implement the following simple recipe:

1. Cluster all the molecules using a diameter merge and a 0.65 threshold.
2. Separate all the molecules in the most populated cluster and assign each of them to an individual bit feature.
3. Collect the remaining leaf clusters and treat them as individual bit features.
4. Pass all the bit features from steps 2 and 3 through a new tree with a threshold of 0.7 (or any desired threshold) and using a tolerance of 0. These steps are represented in Figure 6.

**Figure 6:**
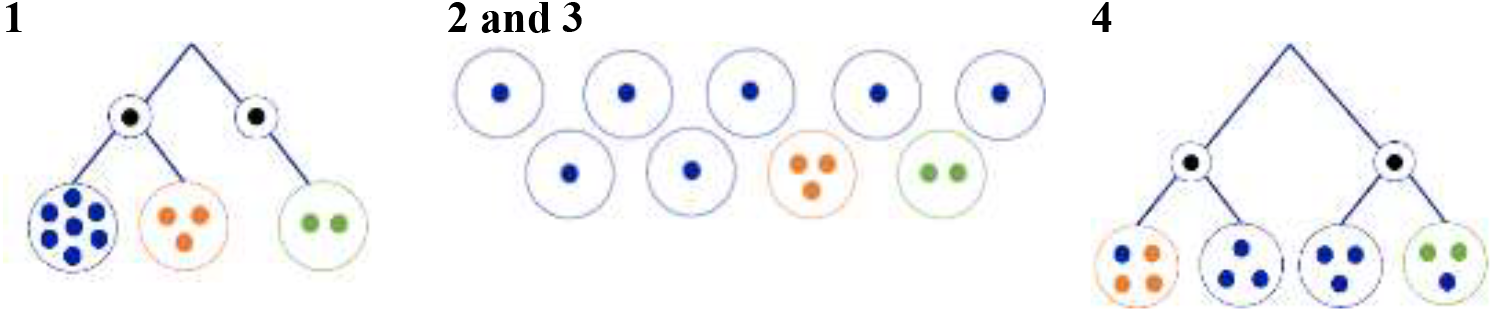
Graphical representation of the proposed composite strategy.

As shown in Fig. 7, this method provides an extremely clean partition of the data, with a uniform distribution of populations and very low scaffold counts per cluster in comparison to the other methodologies. The separation on the t-SNE (Fig 7B) is on par with the one shown in Figures 4H and 4J. As far as a simple, stand-alone procedure, this combination is quite robust. For example, when extracting the BitFeatures of an initial tree and clustering them in a new tree we avoid the biased tree structure that could remain after the pruning, and also do not limit the merging of cluster/molecules that did not encounter each other to the top clusters. Also, the populations of the top clusters are fine-tunable with a second threshold. Although we are creating a second tree, the time overhead is minimal: the second tree will be built with fewer iterations since the number of leaves from the first tree is much lower than the total number of molecules. In this particular example, gathering of the BitFeature and fitting the second tree only took 4.65 seconds.

**Figure 7:**
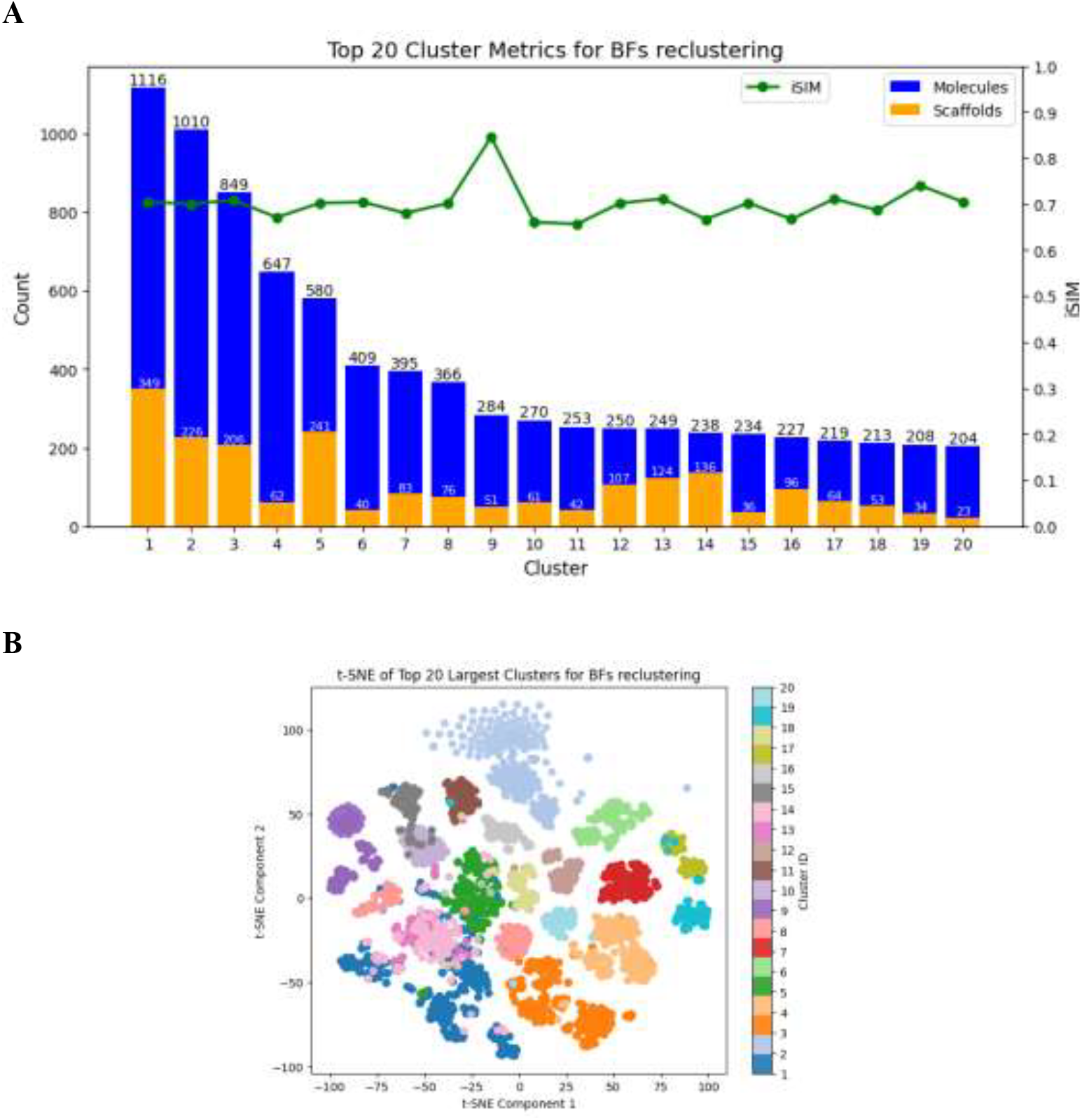
Analysis of the top 20 most populated clusters from the ChEMBL33 natural products set using the composite strategy (Diameter + BitFeature reclustering).

## 5. CONCLUSIONS

This paper introduces several new functionalities to the BitBIRCH algorithm, and effectively serves as the official release of the BitBIRCH software package. This was motivated by the necessity to overcome some of the limitations of algorithms like Taylor-Butina and the original radial version of BitBIRCH, in particular, the formation of a big cluster with an overly-diverse set of scaffolds. Additionally, we want to give the users more control over the clustering process and prepare and maintain a package that is as versatile as possible.

The first BitBIRCH implementation only had a radial merge criterion, but this can be more permissive than desired in some applications. In the newly introduced *diameter* criterion, the comparisons between all the molecules in the set are taken into consideration, resulting in much tighter clusters. Since this can still lead to a relatively big top cluster, we propose a *pruning* alternative that allows removing the biggest cluster from the tree, updates the latter, then it reinserts those molecules. In order to limit the “contamination” with the previously formed clusters, we introduced a *tolerance* check that guaranties that tight clusters are not affected by the reinserted compounds. This improves the previous criteria, significantly reducing the size of the most populated cluster after pairing it with the pruning strategy. Finally, inspired by the original BIRCH implementation, a *reassign* option was included that allows to “relax” the molecules in the top clusters by comparing them to the cluster centroids.

We showcased the flexibility of this package by implementing a composite strategy based on the diameter and tolerance options. This approach uses two trees and performs the second clustering step over the BitFeatures of the first one, while using a tolerance penalty. This resulted in a very robust strategy, with excellent molecule and scaffold counts per cluster. More importantly, this opens the door for clustering practitioners to design their own protocols and pipelines based on the BitBIRCH algorithm.

## Acknowledgements

We thank support from the National Institute of General Medical Sciences and the National Institutes of Health under award number R35GM150620.

## Data and Software Availability Statement

The BitBIRCH code is publicly available at: https://github.com/mqcomplab/bitbirch.

## Notes

### Competing Interest Statement

The authors have declared no competing interest.

## References

(1) Eckert, H.; Bajorath, J. Molecular Similarity Analysis in Virtual Screening: Foundations, Limitations and Novel Approaches. Drug Discov Today 2007, 12 (5–6), 225–233. 10.1016/j.drudis.2007.01.011.

(2) Johnson, M. A.; Maggiora, G. M. Concepts and Applications of Molecular Similarity, 1st ed.; Wiley-Interscience, 1990.

(3) Martin, Y. C.; Kofron, J. L.; Traphagen, L. M. Do Structurally Similar Molecules Have Similar Biological Activity? J Med Chem 2002, 45 (19), 4350–4358. 10.1021/jm020155c.

(4) Bajusz, D.; Rácz, A.; Héberger, K. Why Is Tanimoto Index an Appropriate Choice for Fingerprint-Based Similarity Calculations? J Cheminform 2015, 7 (1), 20. 10.1186/s13321-015-0069-3.

(5) Rogers, D. J.; Tanimoto, T. T. A Computer Program for Classifying Plants. Science (1979) 1960, 132 (3434), 1115–1118. 10.1126/science.132.3434.1115.

(6) Bajusz, D.; Rácz, A.; Héberger, K. Fingerprints, and Other Molecular Descriptions for Database Analysis and Searching - Chemical Data Formats, Fingerprints, and Other Molecular Descriptions for Database Analysis and Searching. In Comprehensive Medicinal Chemistry II; Elsevier, 2017; Vol. 3.

(7) Bajusz, D.; Rácz, A.; Héberger, K. Chemical Data Formats, Fingerprints, and Other Molecular Descriptions for Database Analysis and Searching. Comprehensive Medicinal Chemistry III 2017, 3–8, 329–378. 10.1016/B978-0-12-409547-2.12345-5.

(8) Dunn, T. B.; López-López, E.; Kim, T. D.; Medina-Franco, J. L.; Miranda-Quintana, R. A. Exploring Activity Landscapes with Extended Similarity: Is Tanimoto Enough? Mol Inform 2023, 42 (7). 10.1002/minf.202300056.

(9) Anastasiu, D. C.; Karypis, G. Efficient Identification of Tanimoto Nearest Neighbors. Int J Data Sci Anal 2017, 4 (3), 153–172. 10.1007/s41060-017-0064-z.

(10) Sridhar, D.; Fakhraei, S.; Getoor, L. A Probabilistic Approach for Collective Similarity-Based Drug–Drug Interaction Prediction. Bioinformatics 2016, 32 (20), 3175–3182. 10.1093/bioinformatics/btw342.

(11) Ding, H.; Takigawa, I.; Mamitsuka, H.; Zhu, S. Similarity-Based Machine Learning Methods for Predicting Drug–Target Interactions: A Brief Review. Brief Bioinform 2014, 15 (5), 734–747. 10.1093/BIB/BBT056.

(12) Maggiora, G.; Vogt, M.; Stumpfe, D.; Bajorath, J. Molecular Similarity in Medicinal Chemistry. J Med Chem 2014, 57 (8), 3186–3204. 10.1021/jm401411z.

(13) Buonfiglio, R.; Engkvist, O.; Várkonyi, P.; Henz, A.; Vikeved, E.; Backlund, A.; Kogej, T. Investigating Pharmacological Similarity by Charting Chemical Space. J Chem Inf Model 2015, 55 (11), 2375–2390. 10.1021/acs.jcim.5b00375.

(14) Oyewole, G. J.; Thopil, G. A. Data Clustering: Application and Trends. Artif Intell Rev 2023, 56 (7), 6439–6475. 10.1007/s10462-022-10325-y.

(15) Jain, A. K.; Murty, M. N.; Flynn, P. J. Data Clustering. ACM Comput Surv 1999, 31 (3), 264–323. 10.1145/331499.331504.

(16) Downs, G. M.; Willett, P.; Fisanick, W. Similarity Searching and Clustering of Chemical-Structure Databases Using Molecular Property Data. J Chem Inf Comput Sci 1994, 34 (5), 1094–1102. 10.1021/ci00021a011.

(17) Eljack, F.; Eden, M.; Kazantzi, V.; El-Halwagi, M. Property Clustering and Group Contribution for Process and Molecular Design. Computer Aided Chemical Engineering 2006, 21 (C), 907–912. 10.1016/S1570-7946(06)80161-6.

(18) Beroza, P.; Crawford, J. J.; Ganichkin, O.; Gendelev, L.; Harris, S. F.; Klein, R.; Miu, A.; Steinbacher, S.; Klingler, F.-M.; Lemmen, C. Chemical Space Docking Enables Large-Scale Structure-Based Virtual Screening to Discover ROCK1 Kinase Inhibitors. Nat Commun 2022, 13 (1), 6447. 10.1038/s41467-022-33981-8.

(19) Domingo-Fernández, D.; Gadiya, Y.; Mubeen, S.; Healey, D.; Norman, B. H.; Colluru, V. Exploring the Known Chemical Space of the Plant Kingdom: Insights into Taxonomic Patterns, Knowledge Gaps, and Bioactive Regions. J Cheminform 2023, 15 (1), 107. 10.1186/s13321-023-00778-w.

(20) Hadipour, H.; Liu, C.; Davis, R.; Cardona, S. T.; Hu, P. Deep Clustering of Small Molecules at Large-Scale via Variational Autoencoder Embedding and K-Means. BMC Bioinformatics 2022, 23 (S4), 132. 10.1186/s12859-022-04667-1.

(21) Butina, D. Unsupervised Data Base Clustering Based on Daylight’s Fingerprint and Tanimoto Similarity: A Fast and Automated Way To Cluster Small and Large Data Sets. J Chem Inf Comput Sci 1999, 39 (4), 747–750. 10.1021/ci9803381.

(22) Taylor, R. Simulation Analysis of Experimental Design Strategies for Screening Random Compounds as Potential New Drugs and Agrochemicals. J. Chetn. Inf. Comput. Sci 1995, 35, 59–67.

(23) Lopez-Perez, K.; Jung, V.; Chen, L.; Huddleston, K.; Miranda-Quintana, R. A. BitBIRCH: Efficient Clustering of Large Molecular Libraries. Digital Discovery 2025. 10.1039/D5DD00030K.

(24) Daura, X.; Van Gunsteren, W. F.; Mark, A. E. Folding-Unfolding Thermodynamics of a-Heptapeptide From Equilibrium Simulations. 10.1002/(SICI)1097-0134(19990215)34:3.

(25) Daura, X.; Gademann, K.; Jaun, B.; Seebach, D.; van Gunsteren, W. F.; Mark, A. E.; Rigault, A.; Siegel, J.; Harrowfield, J.; Chevrier, B.; Moras, D.; Lehn, J.; Garrett, M.; Koert, U.; Meyer, D.; Fischer, J. Peptide Folding: When Simulation Meets Experiment. Angew. Chem. Int. Ed. Engl 1998, 31, 1387–1404. 10.1002/(SICI)1521-3773(19990115)38:1/2.

(26) Chen, L.; Smith, M.; Roe, D. R.; Miranda-Quintana, R. A. Extended Quality (EQual): Radial Threshold Clustering Based on n-Ary Similarity. bioRxiv 2024, 2024.12.05.627001. 10.1101/2024.12.05.627001.

(27) López-Pérez, K.; Kim, T. D.; Miranda-Quintana, R. A. ISIM: Instant Similarity. Digital Discovery 2024, 3 (6), 1160–1171. 10.1039/D4DD00041B.

(28) Zdrazil, B.; Felix, E.; Hunter, F.; Manners, E. J.; Blackshaw, J.; Corbett, S.; de Veij, M.; Ioannidis, H.; Lopez, D. M.; Mosquera, J. F.; Magarinos, M. P.; Bosc, N.; Arcila, R.; Kizilören, T.; Gaulton, A.; Bento, A. P.; Adasme, M. F.; Monecke, P.; Landrum, G. A.; Leach, A. R. The ChEMBL Database in 2023: A Drug Discovery Platform Spanning Multiple Bioactivity Data Types and Time Periods. Nucleic Acids Res 2024, 52 (D1), D1180–D1192. 10.1093/nar/gkad1004.

(29) Landrum, G. RDKit: Open-source cheminformatics. https://www.rdkit.org. https://www.rdkit.org (accessed 2025-02-16).

(30) Bemis, G. W.; Murcko, M. A. The Properties of Known Drugs. 1. Molecular Frameworks. J Med Chem 1996, 39 (15), 2887–2893. 10.1021/JM9602928/ASSET/IMAGES/LARGE/JM9602928FB10A.JPEG.

